# Molecular features similarities between SARS-CoV-2, SARS, MERS and key human genes could favour the viral infections and trigger collateral effects

**DOI:** 10.1101/2020.06.23.167072

**Authors:** Lucas L. Maldonado, Laura Kamenetzky

## Abstract

In December 2019 rising pneumonia cases caused by a novel β-coronavirus (SARS-CoV-2) occurred in Wuhan, China, which has rapidly spread worldwide causing thousands of deaths. The WHO declared the SARS-CoV-2 outbreak as a public health emergency of international concern therefore several scientists are dedicated to the study of the new virus. Since human viruses have codon usage biases that match highly expressed proteins in the tissues they infect and depend on host cell machinery for replication and co-evolution, we selected the genes that are highly expressed in the tissue of human lungs to perform computational studies that permit to compare their molecular features with SARS, SARS-CoV-2 and MERS genes. In our studies, we analysed 91 molecular features for 339 viral genes and 463 human genes that consisted of 677873 codon positions. Hereby, we found that A/T bias in viral genes could propitiate the viral infection favoured by a host dependant specialization using the host cell machinery of only some genes. The envelope protein E, the membrane glycoprotein M and ORF7 could have been further benefited by a high rate of A/T in the third codon position. Thereby, the mistranslation or de-regulation of protein synthesis could produce collateral effects, as a consequence of viral occupancy of the host translation machinery due tomolecular similarities with viral genes. Furthermore, we provided a list of candidate human genes whose molecular features match those of SARS-CoV-2, SARSand MERS genes, which should be considered to be incorporated into genetic population studies to evaluate thesusceptibility to respiratory viral infections caused by these viruses. The results presented here, settle the basis for further research in the field of human genetics associated with the new viral infection, COVID-19, caused by SARS-CoV-2 and for the development of antiviral preventive methods.

## 1. Introduction

Since its initial outbreak at Huanan Seafood Wholesale Market in Wuhan, China, in late 2019, COVID-19 has affected more than 4 million people and caused more than 300 thousand deaths all around the world. Thereafter, scientists are focused not only on studying the biology and dissemination of COVID-19 to control the transmission and design proper diagnostic tools and treatments, but also theyare racing to design a vaccine that could prevent the infection caused by the coronavirus SARS-CoV-2. This virus belongs to the Betacoronavirus(β-coronavirus) of the *Coronaviridae*family, which is also composed of three more genera: Alphacoronavirus(αCoV), Gammacoronavirus(γCoV) and Deltacoronavirus(δCoV) (Chen et al., 2020a). Viruses from this family possess a single-stranded, positive-sense RNA and thegenome ranges from 26 to 32 kb (Su et al., 2016).

Coronaviruses have been identified in several host species including humans, bats, civets, mice, dogs, cats, cows and camels (Cavanagh, 2007; Clark, 1993; Wang et al., 2006; Zhou et al., 2018). Since severe acute respiratory syndrome (SARS), caused by the coronavirus SARS-CoV, emerged in southern China in 2002(Peiris et al., 2004), several studies tracing the transmission and possible reservoirs for viruses have been performed. In early 2007, it had already been warned that bats were a natural reservoir for an increasing number of emerging zoonotic virusesas well as for a large number of viruses that have a close genetic relationship with the coronavirusesthat causethe severe acute respiratory syndrome. Furthermore, it was warned that these viruses possess more risk than other pathogens for disease emergence in human and domestic mammals because of their higher mutation rates(Wang et al., 2006). Moreover, legal and illegal trading of wildlife animals propitiates the environment for cross-species virus transmission contributing to the rapid spread of the viral infections around the world(Wang et al., 2006; Wong et al., 2019) as already occurred with SARS and the Middle East respiratory syndrome (MERS) (Zaki et al., 2012). Both viruses have likely originated in bats and are genetically diverse coronaviruses (Cui et al., 2019). Currently, the outbreak of an atypical pneumonia caused by the novel coronavirus SARS-CoV-2 appears to have also startedfrom a zoonotic and a cross-species virus transmission at a market in Wuhan including bats and pangolins, where animals were kept together and the meat was sold(Chan et al., 2020).

Due to viruses replicate exclusively inside of living cells and depend exclusively on the protein synthesis and chaperones machinery of their host, we speculated that the primary structure of viral genes might be determined by the same forces that shape the codon usage in the hosts’ genes. Thereby, viral molecular patterns and codon usage preferences would be a reflection of the host machinery. Codon pair bias and dinucleotide preferences of viruses have been suggestedas the main factors that reflect the codon usage of their hosts. Indeed, virus attenuation by codon pair deoptimization is used as an efficacious attenuation method of various small RNA viruses and has resulted in the generation of superior experimental live virus vaccines (Coleman et al., 2008; Mueller et al., 2010; Nouën et al., 2014; Shen et al., 2015; Wang et al., 2015; Yang et al., 2013).

In order to contribute to solving the sanitary emergence, here we provide a thorough and comprehensive analysis that could help to understand the viability of the virus as well as the susceptibility of the human host to the viral infection based on the molecular patterns of their genes. Therefore, the main goals of ourwork wereto study the molecular and evolutionary aspects of the human coronaviruses SARS-CoV-2, SARS and MERS andto determine the level of similarity of the codon usage and molecular features betweenthe genes of human coronaviruses and the human genesin order to identify the factors that are responsible for the codons selection in the viruses. Moreover, we proposed to identify the essential viral genes for viral replication andhumangenes whosetranslation machinery is involved in propitiating the system for viral replicationin orderto determine whether the genetic population variability could be involved in modelling the gene features andtherefore contributing to the human susceptibility to viral infections.

## 2. Methods

Up to late April, a total of□500 SARS-CoV-2 β-coronavirus genome became available. The total available sequences of β-coronavirus were downloaded from the NCBI (https://www.ncbi.nlm.nih.gov/labs/virus/vssi/#/) including the reference genomes of MERS (NC_019843), SARS (NC_004718) and SARS-CoV-2 (NC_045512) and were classified according to their host. Different SARS-CoV-2 isolates from different countries were pre-analysed but only reference genomes were retaineddue to the low variability of the data. The genomes qualitywas assessed and the genomes containing more than 10 gaps were discarded. CDS of representative viruses fromthe previous classification were selected and analysed. Since human viruses have codon usage biases (CUB) that match highly expressed proteins in the tissues they infect (Miller et al., 2017) we selected 463 highly expressed human genes in lungs tissues according to the fold-change between the expression level in lung and the tissue with second-highest expression level according to the”Human Protein Atlas” (https://www.proteinatlas.org/humanproteome/tissue/lung). We considered valid CDS when they started with an ATG codon, ended with an in-frame stop codon, and had no undetermined nucleotides nor internal stop codons. The accession numbers of the sequences that were used here can be found in Supplementary file 1.

The CUB analyses were performed with CodonW 1.4.4 (J Peden, http://codonw.sourceforge.net/). The total GC content of the CDS as well as the GC content of the first (P1), second (P2), and third (P3) codon positions were calculated using custom PERL scripts. To correct the inequality composition at the third codon position (Sueoka, 1988), the three stop codons (UAA, UAG, and UGA) were excluded in the calculation of P3, and the two single codons for methionine (AUG) and tryptophan (UGG) were excluded from P1, P2, and P3.

### 2.1. Codon usage indices

The following codon indices were calculated: relative synonymous codon usage (RSCU) (Sharp and Li, 1987), the effective number of codons (ENc) (Wright, 1990), codon adaptation index (CAI) (Lee et al., 2010; Sharp and Li, 1987), codon bias index (CBI) (Bennetzen and Hall, 1982), the optimal frequency of codons (Fop) (Ikemura, 1981), General Average Hydropathicity (GRAVY) (Sharp and Li, 1987), aromaticity (Aromo) (Lobry and Gautier, 1994) and GC-content at the first, second and third codon positions (GC1, GC2 and GC3), frequency of either a G or C at the third codon position of synonymous codons (GC3s), the average of GC1 and GC2 (GC12) and Translational selection (TrS2).

ENc indicates the degree of codon bias for individual genes. Over a range of values from 20 to 61, lower values indicate higher codon bias, while ENc equal to 61 means that all codons are used with equal probability (Novembre, 2002; Wright, 1990).

CAI values measure the extent of bias toward preferred codons in highly expressed genes. CAI values range between 0 and 1.0, with higher CAI values indicating higher expression and higher CUB (Lee et al., 2010; Sharp and Li, 1987) under the assumption that translational selection would optimize gene sequences according to their expression levels.

CBI is another measure of directional codon bias, based on the degree of preferred codons used in a gene, like to the frequency of optimal codons. It measures the extent to which a gene uses a subset of optimal codons. In genes with extreme codon bias, CBI will be equal to 1, whereas in genes with random codon usage the CBI values will be equal to 0 (Bennetzen and Hall, 1982).

Fop is a species-specific measure of bias towards particular codons that appear to be translationally optimal in particular species. It can be calculated as the ratio between the frequency of optimal codons and the total number of synonymous codons. Its values range from 0 if a gene contains no optimal codons to 1 if a gene is entirely composed of optimal codons (Ikemura, 1981). The determination of optimal codons was carried out based on the axis 1 ordination, the top and bottom 5% of genes were regarded as the high and low bias datasets, respectively. Codon usage in the two data sets was compared using chi-square tests, with the sequential Bonferroni correction to assess significance according to Peden(Peden, 1999). Optimal codons were defined as those that are used at significantly higher frequencies (p-value < 0.01) in highly expressed genes compared with the frequencies in genes expressed at low levels.

GRAVY values were calculated as a sum of the hydropathy values of all the amino acids encoded by the codons in the gene product divided by the total number of residues in the sequence of the protein. The more negative the GRAVY value, the more hydrophilic the protein is, whereas while the more positive the GRAVY value, the more hydrophobic the protein (Sharp and Li, 1987).

Aromo values denote the frequency of aromatic amino acids (Phe, Tyr, Trp) encoded by the codons in the gene product. (Lobry and Gautier, 1994).

TrS2 estimates the codon-anticodon interaction efficiency revealing bias in favour of optimal codon-anticodon energy and represents the translational efficiency of a gene. TrS2 value > 0.5 shows bias in favour of translational selection according to Gouy and Gautier (Gouy and Gautier, 1982; Uddin et al., 2017; Uddin and Chakraborty, 2018).

### 2.2. Codon Pair Score and Codon Pair Bias

The determination of codon pair biases in coding sequences was performed using CPBias (https://rdrr.io/github/alex-sbu/CPBias/) developed in R. as described by Coleman et al.(Coleman et al., 2008). The CPS is defined as the natural logarithm of the ratio of the observed over the expected number of occurrences of a particular codon pair in all protein-coding sequences of a species. The CPB was used as an index and also to determine the bias in CPS among the virus and host genes. The expected number of codon pair occurrences estimates the number of codon pairs to be present if there is no association between the codons that form the codon pair. It is also calculated to be independent of codon bias and amino acid frequency (Coleman et al., 2008). A negative CPS value means that a particular codon pair is underrepresented, whereas a positive CPS value indicates that a particular codon pair is overrepresented in the analysed protein-coding sequences. Codon pairs that are equally under- or overrepresented have a CPS equidistant from 0. We calculated CPS for each of the 3,721 possible codon pairs (61 x 61 codons).

### 2.3. ENc-plots

The ENc-plot was used to analyse the influence of base the composition on the codon usage in a genome (Hartl et al., 1994). The ENc values were plotted against GC3s values and a standard curve was generated to show the functional relationship between ENc and GC3s values under mutational bias rather than selection pressure. In genes where codon choice is constrained only by a G+C mutational bias, the predicted ENc values will lie on or close to the GC3s standard curve. However, the presence of other factors, such as selection effects, causes the values to deviate considerably from the expected GC3s curve. The values of ENc range from 20 (when only one codon is used per amino acid) to 61 (when all codons are used with equal probability). The predicted values of ENc were calculated according to Hartl*et al*.(Hartl et al., 1994).

### 2.4. Clustering analysis

A total of 91variables of codon usage of viruses and human genes, including the gene composition, RSCU frequencies and the indices described in M&M section, were integrated into an input matrix to feed the clustering algorithm. The analysed variables can be found in supplementary file 1. Hierarchical clustering using Euclidean distances was performed. clValid package clustering algorithm was used to choose and validate the best clustering method. Flexible Procedures for Clustering (fpc:https://cran.r-project.org/web/packages/fpc/index.html) was used for bootstrapping (n=100) and evaluate the cluster stability. Clustering method was used to establish genes groups from codon usage similarities among virus and human genes.

### 2.5. Principal component analysis (PCA)

Principal component analysis (PCA) was used to evaluate the codon usage variation among genes as the multivariate statistical method. The axes represent and allow to identify the most prominent factors contributing to the variation among the genes. Since there are a total of 59 synonymous codons (including 61 sense codons, minus the unique Met and Trpand stop codons), the degrees of freedom were reduced to 40 at removing variations caused by the unequal usage of amino acids during the correspondence analysis of RSCU (Greenacre, 1984). The data were normalized according to Sharp and Li (Sharp and Li, 1987) in order to define the relative adaptiveness of each codon (Peden, 1999; Suzuki et al., 2005), codon usage indices described above were also included as variables. PCA analyses were performed using “factoextra R package” (https://cloud.r-project.org/web/packages/factoextra/index.html).

### 2.6. Phylogenetic analyses

The DNA genome sequences of all the viruses were aligned usingClustalO v1.2.4(Sievers et al., 2011). PartitionFinder 2 (Lanfear et al., 2016) was used to select the best-fit partitioning schemes and models of evolution for the phylogenetic analysisThe evolutionary model was set on generalised time-reversible substitution model with gamma-distributed rate variation across the sites and a proportion of invariable sites (GTR +G). The final phylogeny was calculated using fasttree(Price et al., 2009). The bootstrap consensus trees inferred from 1000 replicates were retainedin the bootstrap and the final trees were drawn usingFigtree (https://github.com/rambaut/figtree/releases).

## 3. Results

### 3.1. Phylogeny

Up to late April, a total of □ 500 SARS-CoV-2 β-coronavirus genomes became available and the number of available genomes incremented substantially. The total available sequences of β-coronavirus were downloaded from the NCBI. In order to evaluate the variability and to select the best candidates for codon usage and nucleotides content analyses, we performed phylogenetic analyses by implementing the GTR + G model according to partition finder results. Firstly, a phylogenetic tree was constructed (data not shown) using the whole genome sequences of SARS-CoV-2 reported in humans from Spain, USA, Italy, South America, China, Korea Japan and Australia, the references genomes of MERS and SARSand the viruses genomes isolated from bats, pangolins, civets, hedgehogs, *Bos taurus*, and canids (Supplementary file 1). Since all SARS-CoV-2 genome sequences remained together in the same cluster we selected representative viruses genomes randomly from each country and from each host and constructed a final phylogenetic tree (Supplementary file 2). The topology of this tree showed that SARS-CoV-2 samples diverge from a commonnode close to the bat virus (Accession MN996532) and that all of them diverge from a common node very close to pangolin viruses (Accessions MT040333, MT040334, MT040335, MT040336, MT072864). From a distant node that contained the node of SARS-CoV-2 and also clustereda set of bat viruses, SARS virus diverges and grouped within a node together with Civets viruses, but also adjacent and very close to other bats viruses (Accession KY417146, KT444582 and KY417150). MERS viruses grouped in a different node also very close to other cluster containingbat viruses (Accession MF593268 and KC869678). Furthermore, adjacent to this node we found hedgehogs viruses (Accession KC545383, KC545386, MK679660, MK907286, MK907287 and NC_039207).

For further analyses of molecular features and codon usage properties of viral genes, we selected the viruses according to the phylogenetic tree described above. First, we identified the 3 nodes containing the human viruses: SARS-CoV-2, SARS and MERS. Then we selected the closest viruses to each human virus within the nodes and classified them according to their hosts (bats, civets, hedgehogs and pangolins). From the human viruses only the references MERS (NC_019843), SARS (NC_004718) and SARS-CoV-2 (NC_045512) were used. The total selected CDSs comprised104 viral CDS.

### 3.2. Viral gene codon usage patterns

PCA of codon usage and molecular features of viral genesof SARS, MERS, SARS-CoV-2 and related viral genes of the non-human host (bats, civets, hedgehogs and pangolins) were performed in order to characterise the genes and distinguish important gene features among the viral gene families and species. PCA showed that the genes dispersed differentially according to the kind of gene rather than tothe host that the viruses infect. The genesdistribution depending on the host was observedfor only some genes(Figure 1 A and B). Most of the genes belonging to the same gene family overlapped or positioned in a very short ratio from each other. The position for nucleocapsid protein N is shared for all virusesexcept for SARS-CoV-2 that is the most distant from the group followed by the ones of SARS and MERS. Regarding the envelope protein E, the gene of civetsSARS and human SARSoccupied the same positionwhile the gene of human SARS-CoV-2 and MERS distributed distantly from the group and alsofrom each other. The genes that encode for the membrane glycoprotein M also showed a distribution along positive axis 1. However, for thehuman SARS-CoV-2 and hedgehog and bat MERSthese genes distributed away from the genes of pangolin and bat SARS-CoV-2, human MERS and bat, civet and humanSARS toward the inferior left quadrant. The main factors that contributedto the dispersion ofthese 3 genes (nucleocapsid protein N, envelope protein E and membrane glycoprotein M) were CBI, Fop, TrS2, C3, C3s, CpG and GC. In addition, the membrane glycoprotein M seems to be more influenced by the codon frequency of the codon TAC for Tyrosine and CTG for Lysine. Whereas for hedgehogs MERS and human SARS-CoV-2 the codons TAT for Tyrosine and TTA for Lysine as well as the GC bias, among others contributed most. All of them are strongly influenced by the A/T composition in the third codon position. On the other hand, the spike proteins S of all viruses distributed toward positive values of PC2and were also highly influenced by the A/T compositionin the third codon position. All these genes groupedvery closelyexcept for the gene of bats and human MERS. Several ORF proteins occupied the same area overlapping or positioning very close to each other. Other ORF genes such as ORF6, ORF8 and ORF3of the human virus SARS-CoV-2 distributed distantly.

**Figure 1:**
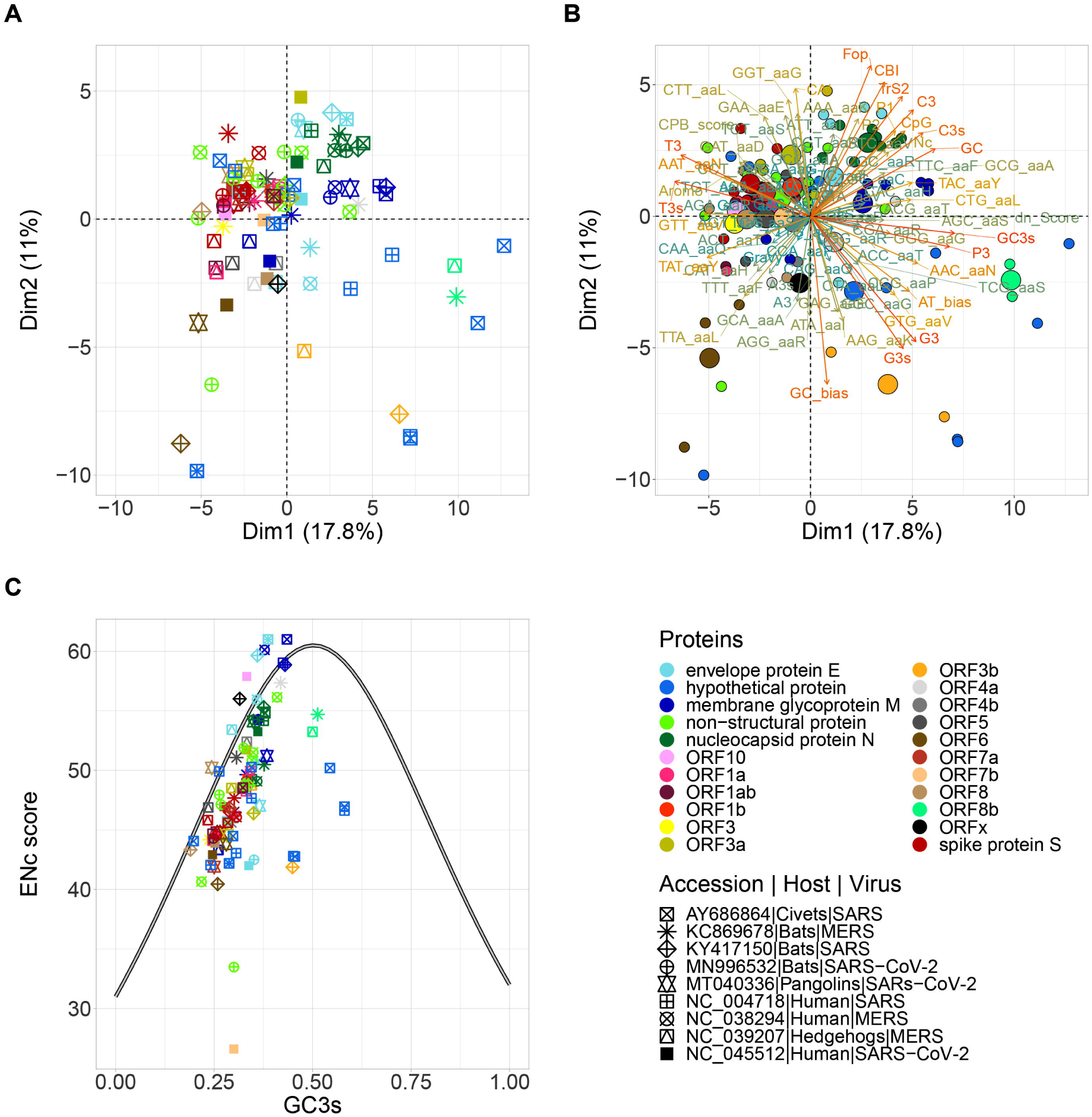
viral genes distribution in PCA plot in the first 2 axes and ENc-GC3s plot of SARS-CoV-2 (NC_045512), SARS (NC_004718) and MERS (NC_038294), and the related virus of non-human hosts: CivetsSARS (AY686864), bats MERS (KC869678), bats SARS (KY417150), bats SARS-CoV-2 (MN996532), pangolins SARS-CoV-2 (MT040336) and hedgehogs MERS (NC_039207). **A)**. Distribution of viral genes in PC1 and PC2. **B)**. Main factors represented by vectors that contribute to the distribution of viral genes in PC1 and PC2. **C)**. Distribution of the effective number of codons (ENc) in relation to the GC3s of viral genes. The standard curve of ENcis indicated in solid line.

The CUBwas estimated based on ENc values. The values of ENc range from 20 (when only one codon is used per amino acid) to 61 (when all codons are used with equal probability). Genes whose ENc values are lower than 50 are considered to have skewed codon usage. All viruses presented a wide range of ENc values (26 <ENc< 58) that varied mainly depending on the type of genes. The highest ENc median was observed for the human MERS (ENc∼50.40), followed by bats MERS (ENc ∼49.64). The hedgehogs MERS showed an ENc median of 47.70, being closer to the values that SARS and SARS-CoV-2 presented. Civets SARS showed an ENc median of 47.76. For bats SARS the ENc median was 47.80 and the ENc median for the human SARS was 47.66. The lowest ENc valueswere observed for the genes of SARS-CoV-2 viruses. Bats SARS-CoV-2 showed a median ENc of 46.39. For pangolins SARS-CoV-2, the ENc median was 44.80 and for the human SARS-CoV-2, the ENc median was 44.34 (Figure 1 C).

Codon bias of viral genes classified bythe type of gene or the gene familyshowed that the envelope protein E presented ENc values that ranged from 42 to 61, being the genes of the human SARS-CoV-2 and bat SARS-CoV-2 the ones that presented the lowest valuesof all the genes. Membrane glycoprotein M genes presented ENc values that ranged from 43.32 to 61.00, and the gene of hedgehogs MERS, the one with the lowest value. The membrane glycoprotein M of Human MERS, Civets, bats and human SARS showed virtually non-biased codon usage. The nucleocapsid protein N showed ENc values that ranged from 49.08 to 55.30 being the human MERS gene, followed by the bats MERSgene, the viral genes that presented the lowest values.

The genes that encode for the spike protein S presented ENc values that ranged from 44.16 to 47.68. The gene of the humanSARS-CoV-2 showed the lowest ENc value followed by the genes of bat and pangolin SARS-CoV-2. ORF genes presented ENc values that ranged from 26.60 to 57.89, being the human SARS-CoV-2 the virus that presented the lowest and the highest ENc value for ORF7 and ORF10 respectively.

### 3.3. CPB analysis between viruses and lung tissue highly expressed genes

Since human viruses have codon usage biases that match highly expressed proteins in the tissues they infect (Miller et al., 2017) we selected the highly expressed genes in lungs tissue to compare the gene molecular features, CUB and CPB of SARS, SARS-CoV-2 and MERS viruses in relation with human genes. Since viruses replicate exclusively inside living cells, many viruses are influenced by host codon pair preferences, being there flection of the CPB or CPS of their hosts. Therefore, CPSof viral genes was evaluated and compared with the CPS of human genesin order todetermine whether viruses that infect humans have similar codon pair preferences to their host. Viral genes showed lower CPS frequencies than human genes. The median CPS for the viral genes was 0.053 for civet SARS, 0.051 for bat SARS, and 0.053 for human SARS. The median CPS that presented the MERS genes was 0.048 for hedgehogMERS, 0.046 for bat MERS and 0.037 for human MERS. SARS-CoV-2 showed median CPS values of 0.064 for bat SARS-CoV-2, 0.065 for pangolins SARS-CoV-2 and 0.061 for human SARS-CoV-2. The median CPS for highly expressed human genes in lungs was 0.11 (Figure 2A). We also calculated the median CPS of viral genes for each gene family. The genes that encode for ORF7b and spike protein S were the genes with the highest median of CPB values (0.081 and 0.061 respectively), followed by ORF3a and nucleocapsid protein N (0.058 and 0.057) indicating that particular codon pairs are overrepresented in these genes. Envelope protein E and the membrane glycoprotein M showed values of 0.048 and 0.044 respectively (Figure 2B).

**Figure 2:**
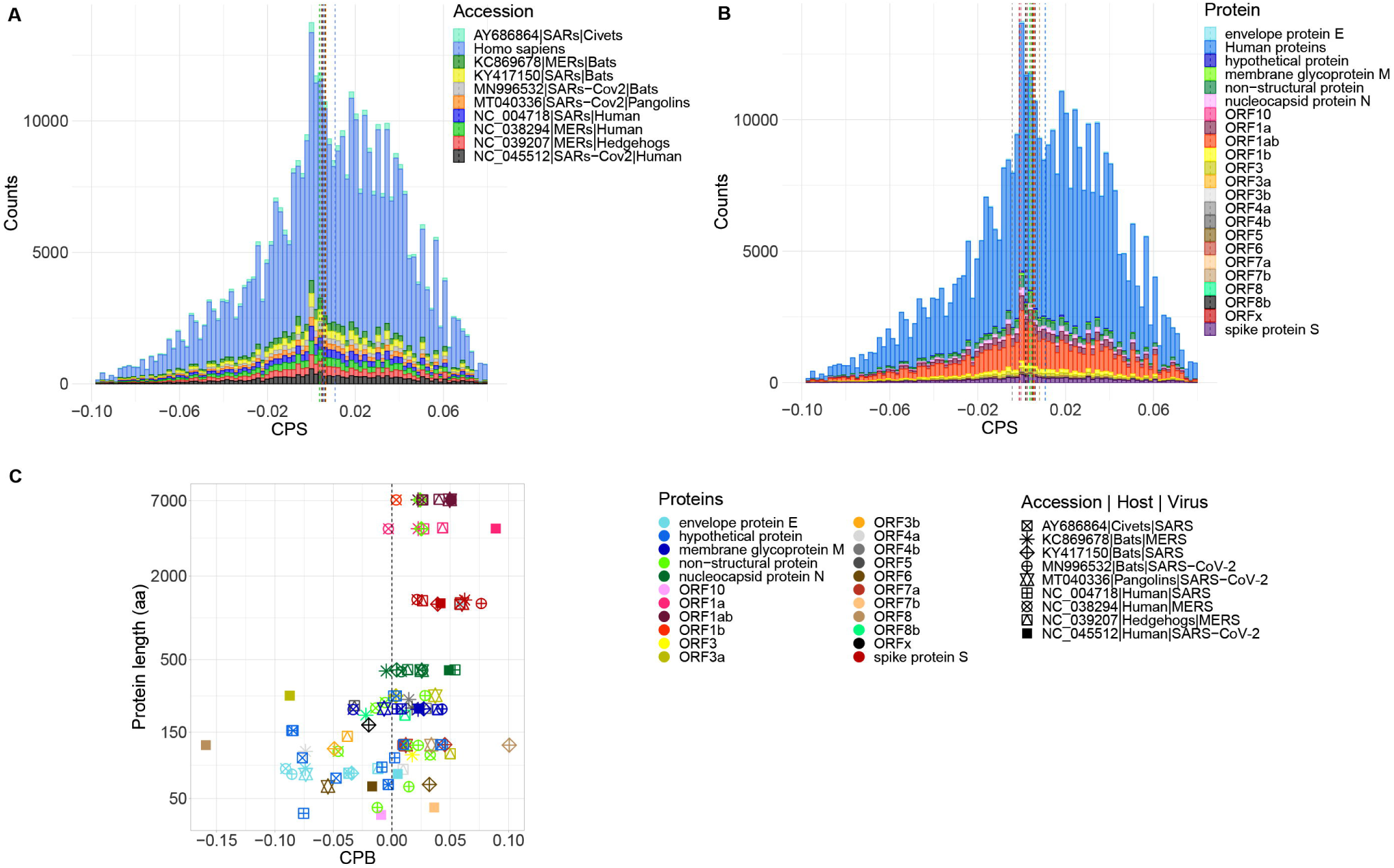
codon pair score of viral genes of SARS-CoV-2 (NC_045512), SARS (NC_004718) and MERS (NC_038294), and the related virus of non-human hosts: civets SARS (AY686864), bats MERS (KC869678), bats SARS (KY417150), bats SARS-CoV-2 (MN996532), pangolins SARS-CoV-2 (MT040336) and hedgehogs MERS (NC_039207) and human genes. **A)**. codon pair frequencies for each virus. **B)**. codon pair frequencies for each gene and classified by type of viral gene. **C)**. Codon pair bias for each viral gene vs protein length.

Furthermore, different CPB among the viral genes families of different virusesspecies that infect different hostswere also observed (Figure 2C). Bat SARS showed the highest CPB for ORF7b (∼0.1) and ORF7a (∼0.048). The human SARS-CoV-2 showed the highest CPB values for ORF1a (0.089), ORF1ab (0.051), nucleocapsid protein N (0.048) and spike protein S (0.042). Envelope protein E and membrane glycoprotein M presented CPB values of 0.0005 and 0.02 respectively. In pangolin SARS-CoV-2 the highest CPB value was for the spike protein S (0.060) followed by ORF3a (0.037) and ORF8 (0.033) while inbat SARS-CoV-2 the highest CPB value was for the spike protein S (0.076) followed by ORF1ab (0.050). The highest CPB values for human MERS genes were observedfor a non-structural protein (0.032) followed by the spike protein S (0.021) and the nucleocapsid protein N (0.008). In bat MERS the highest CPB value was for spike protein S(0.062) followed by ORF1a (0.022) and by the membrane glycoprotein M (0.022). In hedgehog MERS the highest CPB value was for ORF3a (0.049), followed by ORF1a/b (0.042) and the membrane glycoprotein M (0.029). In orderto evaluate the fitness and specialization of the viruses in theirhosts, we compared the CPS of the viral genes derived from the different hosts against the CPS of highly expressed genes in human lungs tissue by performing CPS correlation analysis. The 3721 codon pairs of all viruses were compared with the 3721 codon pairs of human genes. Correlation analyses showed low R values with a not clear dependence on the human host codon pairs (Supplementaryfile3).

### 3.4. Viral genes clustering analysis

Further analysis using hierarchical clustering of viral genes provided additional qualitative information about how similar certain genes are in terms of molecular and codon usage parameters(Supplementaryfile4). All the genes encoding for viral nucleocapsid protein N of all the virusesgrouped together demonstrating a high level of conservation. The human SARS-CoV-2 gene presented a very low CPB value in comparison with the genes of the other viruses.

Envelope protein E genes grouped in different clusters. Thethree SARS viruses (civets, bats and human hosts) grouped with the bat and pangolin SARS-CoV-2 and showed high and similar CPB values. Whereas the human SARS-CoV-2 grouped better with the human and bat MERS, both showinghigh CPB values. Hedgehog envelope protein E gene grouped distantly with ORF and hypothetical proteins. These genes presented a high preference for the codon GCG for Alanine. Furthermore, the codons GGT and CCT were preferable for Glycine and Proline respectively. The hydrophobicity, CAI and Fop values are higher for SARS and SARS-CoV-2 genes.

The membrane glycoprotein M genes also appeared in different clusters. The humanSARS-CoV-2 gene grouped with the human and bat MERS gene. Only the human SARS-CoV-2 gene showed a preference for the codon GCA for Alanine and AGG for Arginine. A preference for the codons CCA, ACT and GTT for Proline, Threonine and Tyrosinewas also observed. However, this was the gene that presented the lowest CPB value. The pangolin and bat SARS-CoV-2 gene grouped with the threeSARS genes in a different clusterand showed a higher CPB. The hedgehogs MERS grouped distantly. The codon CAC was observed more frequently for Histidine in the three SARS viruses. Pangolin SARS-CoV-2 showed a preference for GAC for Aspartate and CCA for Proline whereas the bat SARS-CoV-2 showed a preference for GGA for Glycine and GTA for Valine.

The genes encoding for the spike protein S appeared in different clusters. For the human SARS-CoV-2, this gene grouped alone with the genes that encode for hypothetical and ORF proteins and showed high bias only for the codon TTG for Lysine and GAA for Glutamate. The genes for the rest of the viruses clustered together with other ORF genesand with the membrane glycoprotein M of hedgehogs, although distantly. This analysis also showed that CPB is highly related to the dinucleotide bias. In this cluster the genes that encode for the ORF genes showed low values of CPB, being the lowest of all the clusters. Conversely, the genes that encode for the spike protein S presented high CPB values.

### 3.5. Viral and human genes clustering analysis and CPS correlation analysis

Since virus fitness specialization could be dependent on the translation machinery for only some particular genes, we performed clustering analysis based on the codon usage and molecular features including human genes and the genes of SARS-CoV-2, SARS and MERS(Supplementary file 1). From a total of 463 human genes, 70 genes (15.1%) grouped in clusters together with viral genes. These genes were selectedand extracted to make an illustrating heatmap including both, human and all viral genes (Figure 3). Furthermore, in order to figure out whether the viral genes that composed particular clusters present codon pair frequencies that correlate with the human genes of the same clusters, we evaluated the CPS of both, viral and human genes for each block and the correlation between each other.

**Figure 3:**
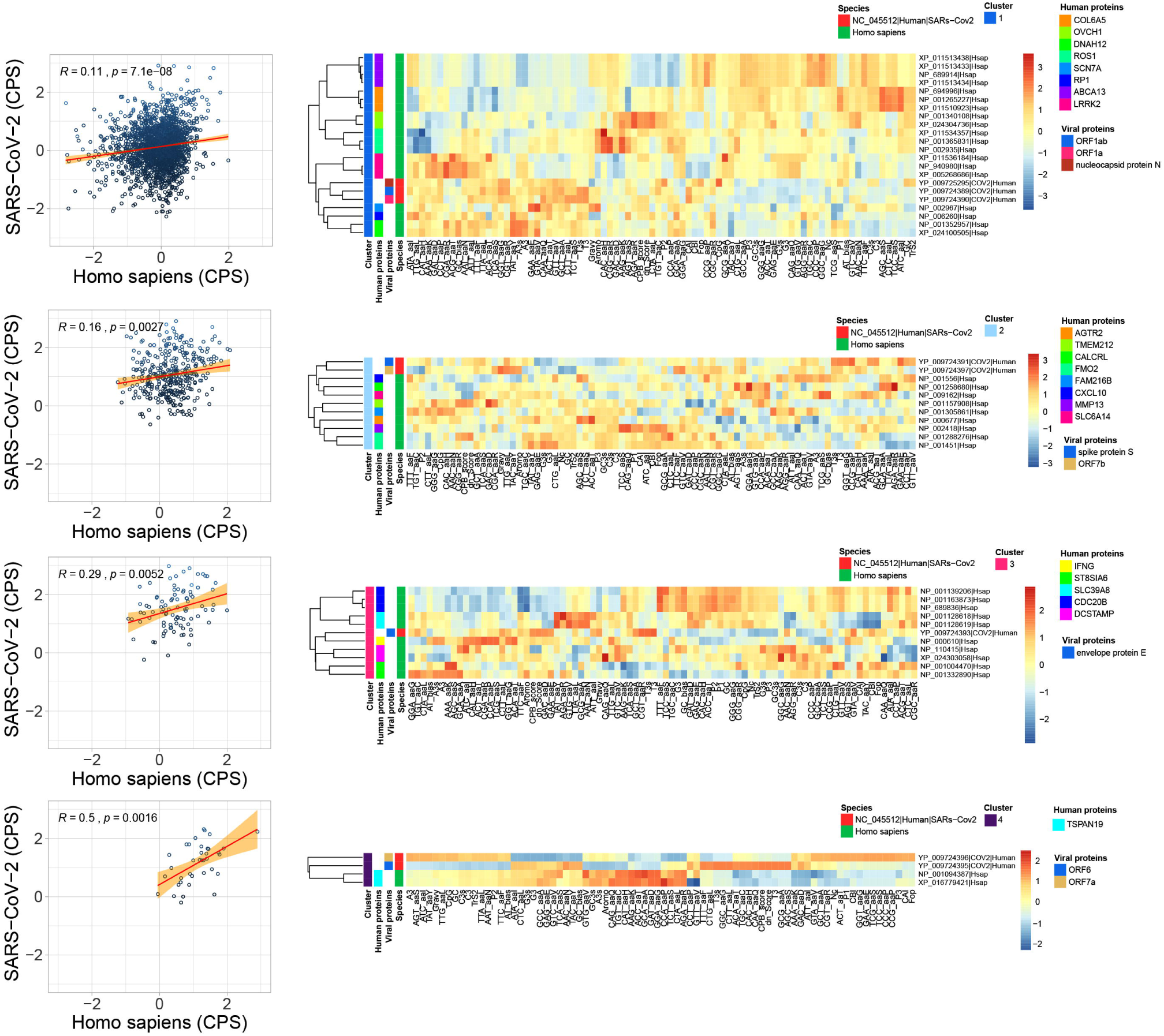
Heatmap of clusters(1 to 4) using a hierarchical method of viral genes forSARS-CoV-2 (NC_045512) of the human host and human genes based on the molecular features. CPB correlation is included in the left for each cluster relating the CPB of human genes (horizontal axis) and CPB of the viral genes (vertical axis).

For SARS-CoV-2, 8 out of 12 genesgrouped with 40 human genes distributed in 4 clusters. The first cluster comprised3 viral genes: the nucleocapsid protein N, ORF1a and ORF1ab together with 19 human genes: COL6A5 (3), OVCH1 (2), DNAH12 (2), ROS1 (3), SCN7A, RP1, ABCA13 (4) and LRRK2 (3) and the CPS correlation was R□0.11 (p-val□7.1×10^−8^). The CPB values for the human and viral genes were −0.04 and 0.15 respectively. The second cluster contained 2 viral genes: the spike protein S and ORF7b together with 9 human genes: AGTR2, TMEM212, CALCRL, FMO2 (2), FAM216B, CXCL10, MMP13 and SLC6A14. For this group, the CPS correlation between viral and human genes was R□0.21 (p-val□0.0027) and the CPB values for the human and viral genes were 0.05 and 0.12 respectively. The third cluster grouped only 1 viral gene, the envelope protein E, with 10 human genes: IFNG, ST8SIA6 (2), SLC39A8 (2), CDC20B (3) and DCSTAMP (2), and the CPS correlation between viral and human genes wereR□0.29 (p-val□0.0052) and the CPB values for the human and the viral genes were 0.25 and 0.16 respectively. In the fourth cluster 2 human genes, TSPAN19 (2) grouped with the viral genes ORF6 and ORF7a and showed a CPS correlation of R□0.5 (p-val□0,0016). The CPB values for the human and viral genes were 0.13 and 0.14 respectively.

For SARS virus (Supplementary file 5), 12 out of 14 genes grouped with 39 human genes distributed in 5 clusters. The first cluster contained 11 human genes: AGTR2, TMEM212, CALCRL, FMO2 (2), FAM216B, CXCL10, MMP13, SLC6A14, and BCL2A1 (2) and 7 viral genes: envelope protein E, hypothetical proteins (5) and membrane glycoprotein M. The CPS correlation for human and viral genes was R□0.28 (p-val□2×10-^16^). The second cluster was composed of 7 human genes: C7orf77, SNTN (2), MUC1 (2) and WIF1 (2). The CPS correlation between human and viral genes was R□0.77 (p-val□0.00078). The third cluster consisted of 19 human genes: COL6A5 (3), OVCH1 (2), DNAH12 (2), ROS1 (3), SCN7A, RP1, ABCA13 (4) and LRRK2 (3) and 2 viral genes: ORF1a and ORF1ab. The CPS correlation between human and viral genes was R□0.31 (p-val□2.2×10-^16^). The fourth cluster was composed of 2 TSPAN19 human genes and one viral hypothetical protein. CPS correlation for this cluster was statistically not significant.

For MERS virus (Supplementary file 6), the 10 genes grouped into 6 clusters together with 65 human genes. The first cluster comprised 19 human genes: COL6A5 (3), OVCH1 (2), DNAH12 (2), ROS1 (3), SCN7A, RP1, ABCA13 (4) and LRRK2 (3) and 2 viral genes: ORF1a and ORF1b. CPS correlation for human and viral genes was R□0.29 (p-val□0.021). The second cluster was composed of 8 human genes: AGTR2, TMEM212, CALCRL, FMO2 (2), CXCL10, MMP13 and SLC6A14 and 1 viral gene that encodes for the spike protein S. The third cluster contained 6 human genes: CLEC12A (3), FAM216B, DNAAF6 and PIH1D3 and 3 viral genes: non-structural protein (2) and the nucleocapsid protein N. The fourth cluster consisted of 7 human genes: C7orf77, SNTN(2), MUC1 (2) and WIF1 (2) and 1 viral gene encoding for a non-structural protein. The fifth cluster was composed of 9 human genes: ST8SIA6 (2), SLC39A8 (2), CDC20B (3) and DCSTAMP (2) and 2 viral genes: non-structural protein and membrane glycoprotein M. The sixth cluster comprised 16 human genes: CLEC6A, IL1RL1 (2), VNN2 (4), RTKN2 (2), SDR16C5 (3), IL18R1 (2), ACADL, and DNAH12 and the viral gene that encodes for envelope protein E.

Furthermore, we observed that 28 out of 70 human genes comprised a core of genes that appeared into the clusters together with the viral genes for the three human viruses (Figure 4 and Table 1). Between MERS and SARS-CoV-2 only 9 human geneswere shared, 7 genes were shared between onlyMERS and SARS and 2 genes were shared between only SARS-CoV-2 and SARS. The 28 human genes that appeared in the clusters together with the genes of the 3 human viruses were retrieved against PheGenI (https://www.ncbi.nlm.nih.gov/gap/phegeni/#pgGAP) and DisGeNET (https://www.disgenet.org/) in order to classify the human diseases associated with the malfunction of the identified genes as an approach for possible human diseases or collateral effects caused by the viral infections (Supplementary file 1). According to PheGenI and DisGeNET,14 out of the 28 genes were associated with 27 diseases. All the diseases appeared in nearly equal proportions. A slightly higher frequency was observed for Nervous System Diseases (12 genes), Neoplasms (12 genes), Respiratory Tract Diseases (11 genes), Pathological Conditions Signs and Symptoms(11 genes), Neonatal Diseases and Abnormalities (11 genes) and Cardiovascular Diseases (10 genes) among others (Figure 4 B).

**Table 1:**
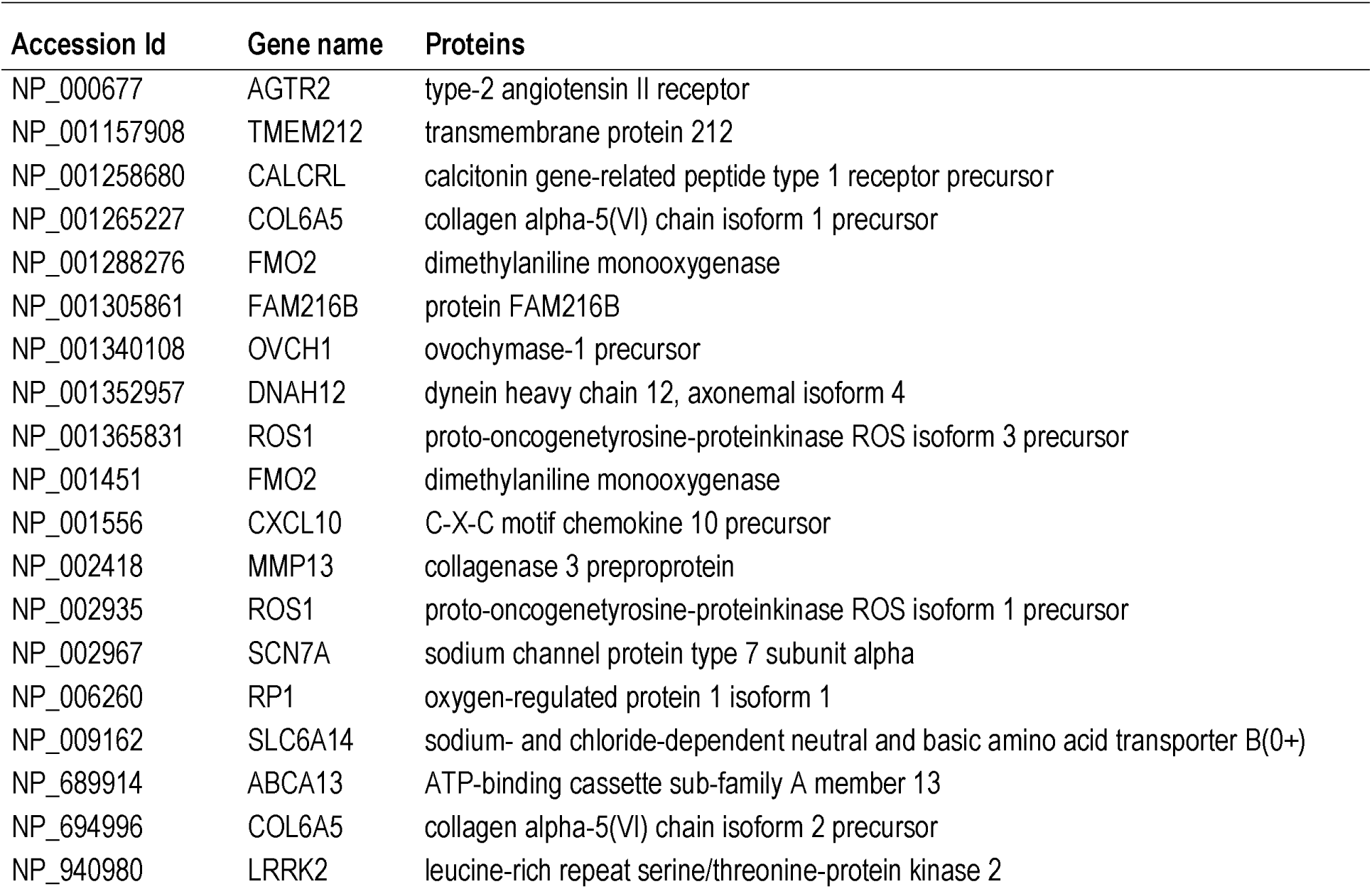
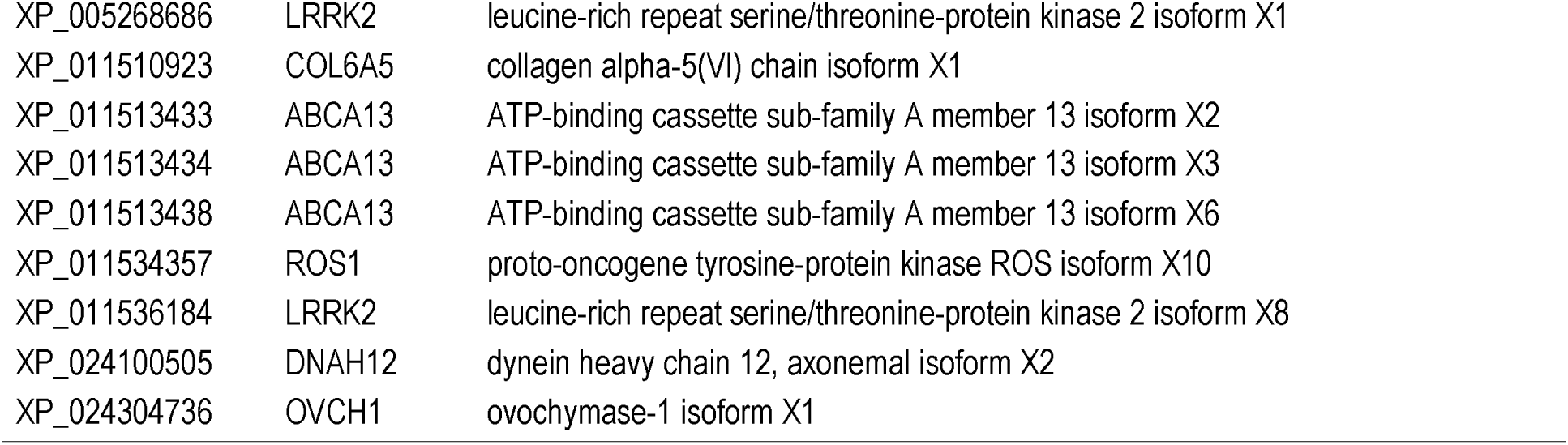
Human genes shared among the clusters of the three coronaviruses SARS-CoV-2, SARS and MERS.

**Figure 4:**
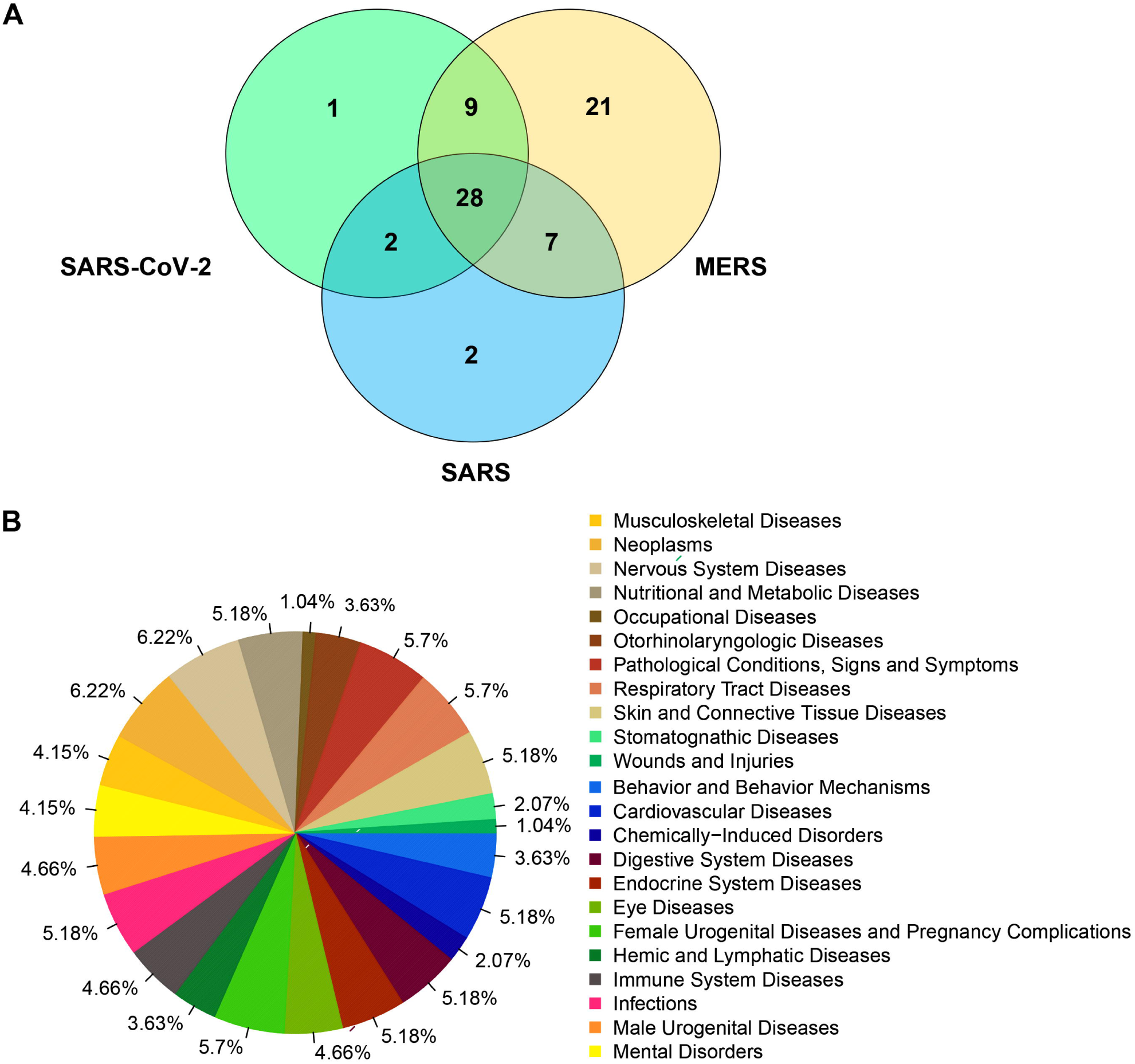
A). Venn diagram representing the number of human genes that clustered together with viral genes for SARS-CoV-2 (NC_045512), SARS (NC_004718) and MERS (NC_038294) based on the molecular features. **B)**. Diseases frequencies associated to human genes grouped with viral genes of SARS-CoV-2, SARS and MERS in the clustering analysis.

## 4. Discussion

In humans, coronaviruses cause mainly respiratory tract infections. Previously, six coronaviruseswere identified as human-susceptible viruses, being theβ-coronaviruses, SARS-CoV and MERS-CoV, responsible for severe and potentially fatal respiratory tract infections (Yin and Wunderink, 2018). In December 2019 rising pneumonia cases caused by a novel β-coronavirus (SARS-CoV-2), occurred in Wuhan, China. The disease was officially named coronavirus disease 2019 (COVID-19). It was found that the genome sequence of SARS-CoV-2 is 96.2% identical to the bat coronavirus RaTG13, whereas it shares 79.5% of identity to SARS-CoV. It has been proposed that SARS-CoV-2 may have originated from bats or unknown intermediate hosts that could involve pangolins, crossing the species barrier into humans. (Guo et al., 2020). Rather, bats are the natural reservoir of a wide variety of coronaviruses, including SARS-CoV-like and MERS-CoV-like viruses (Banerjee et al., 2019; Hampton, 2005; Li et al., 2005).

Up to late April, a total of □ 500 SARS-CoV-2 β-coronavirus genomes became available and this number is continuously increasing at an unprecedented rate. In our studies, we analysed the coronaviruses from different host species and retrieved the phylogeny in order to select the best candidate genomes for further analysis of codon usage and molecular relationship with the human host. As previously reported, we found that for SARS-CoV-2, SARS and MERS, bats seem to be a common natural host or reservoir.

Because viruses do not have tRNAs and rely on host cell machinery for replication, co-evolution between RNA viruses and their hosts’ codon usages have been observed (Franzo et al., 2017; Rahman et al., 2018; Simón et al., 2017). Furthermore, the adaptation of viruses that replicates in multiple hosts should involve a trade-off between precise and functional matching to fit the diverse tRNA pools of multiple hosts (Tian et al., 2018). Conversely, single host viruses are expected to have specialised to match the host tRNA repertoire.

Since human viruses have CUB that matches highly expressed proteins in the tissues they infect (Miller et al., 2017) we selected the highly expressed genes in lung tissues to compare gene molecular features of SARS-CoV-2, SARS and MERS in relation with human genes. In general, in our studies for all the analysed viruseswe found that the total gene repertoire had a similar ENc average that differs only [1 unit with respect to their non-human host they come from, reflecting the molecular features of their original host. Furthermore, as demonstrated in our clustering analysis, codon pair usage seems to be dependent on the dinucleotide bias and the human CPB was higher for human genes than for viruses genes as previously reported (Kames et al., 2020; Kunec and Osterrieder, 2016).

Moreover, our analyses allowed us to distinguish not only the main factors that contribute to the distribution of the genes along the axes in PCA, but also to determine some particular different features among human and non-human viruses in specific genes that could be important for explaining the virus infection evolution. In contrast to SARS-CoV-2 of bats and pangolins, human SARS-CoV-2 exhibited a differential distribution in particular genes that depended mostly on the A/T content in the third codon position, which is in accordance with human SARS-CoV-2 gene composition reports (Alnazawi et al., 2017; Alonso and Diambra, 2020; Kames et al., 2020; Tort et al., 2020).

Important viral genes such as the membrane glycoprotein M, that is involved in the membrane transport of nutrients, the bud release, the formation of the envelope, the virus assembly and in the biosynthesis of new virus particles (Guo et al., 2020), distributed differentially from the non-human viruses indicating that it is highly influenced by A/T content. Surprisingly, this gene was positionednear of the hedgehog’s MERS gene, suggesting similar molecular patterns between two distant viruses.

The envelope protein E, that functions as an ion channel and regulates virion assembly and the immune system of the host (Guo et al., 2020; Yin and Wunderink, 2018), showed the same tendency toward A/T ending codons for the human viruses SARS-CoV-2 and MERS. Human genes using A/T ending codons is also a common feature in several human genes since they present a wide rate of GC content (27-97%) (Bernardi, 2015; D’Onofrio et al., 1991). Both, membrane glycoprotein M and envelope protein E genes have a higher CUB in comparison with human MERS and SARS which is not in accordance with the trade-off theory that postulates that cross-species virus transmission demands relaxing the codon usage pattern (Kunec and Osterrieder, 2016). However, this phenomenon could be explainedby a selection pressure in favour of the virus replication in the new host or due to the recent cross-species virus transmission as we know it occurred. If this is the case, the analysis of newisolates from infectedhumans should tend to show incremented ENc values for the envelope protein E and membrane glycoprotein M genes.

Nevertheless, only the envelope protein E clustered together with human genes, demonstrating similar molecular patterns that could mean an advantage for virus replication in humans facilitating the virion assembly and the regulation of the immune system of the host. Furthermore, positive CPB and an incremented CPS correlation for the cluster that grouped the envelope protein E with human genes supports the hypothesis of a facilitated translation depending on codon usage and codon pairs. Similar patterns were observed for ORF6 and ORF8genes, which are involved in the viral pathogenesis, apoptosis induction and inflammatory responses in the host (Chen et al., 2020b; Diemer et al., 2010). These genes grouped with human genes in different clusters and showed also and incremented CPS correlation.

Studies in different viruses species have reported high conservation of the genes that encode for the nucleocapsid protein N among virus families (HORNE, 2013; Kunec and Osterrieder, 2016; Masters, 2019; Nathan and Scobell, 2012; Parker and Masters, 1990). Therefore, it is expected that codon usage and molecular patterns present similar features as observed in our studies. Nevertheless, SARS-CoV-2 nucleocapsid gene tends to distribute slightly far from the rest of viruses’ capsids genes and toward a higher A/T content in PCA. Both, the human SARS and SARS-CoV-2 nucleocapsid protein N present high CPB suggesting a specialization acquirement in the human host.

Two viral genes that also present high CPB are ORF1a/b, that encodes for the replicase complex (polyproteins pp1a and pp1ab) and the Spike protein S that participates in the early viral infection by attaching to the host receptor ACE2 and mediating the internalization of the virus (Guo et al., 2020). In our studies, ORF1a/b grouped with the gene that encodes for the nucleocapsid protein N, indicating that their molecular features are highly conserved and are also presentin several human genes. This result is in concordance with previous works that proposed these genes as candidates for deoptimization for the design of attenuated vaccines due to their high positive CPB values (Kames et al., 2020). Instead, the gene that encodes for the spike protein S, grouped with ORF7 (involved in viral pathogenesis and apoptosis induction) that also presents high and similar positive CPB values. For all of them, a higher rate of A/T composition in the third codon position was observed. Changes in the third position produce synonymous substitutions that could have conducted to a codon optimization in human cells using the host machinery that translates only genes whose molecular features match the viral needs. Some viral genes seem to have been favoured for an increased viral replication in humans and optimized by using or mimicking some particular molecular patterns of human genes. But only some genes, such as the envelope E, the ORF 6 and 8, could be the key for an exacerbated viral pathogenesis. Furthermore, because of these molecular and codon usage similarities between some highly expressed human genes and viral genes that occupy the same clusters, the translation machinery of the host could propitiate the translation of viral genes to the detriment of human gene expression in lung tissues. Indeed, mistranslation or de-regulation of protein synthesis has been reported as a consequence of tRNA miss-modification and imbalanced tRNA expression, causing diseases(Lant et al., 2019). Recent studies have also proposed that an unbalance in the tRNAs pools of the infected cells could occur and would explain the collateral effects observed in some viral infections (Alonso and Diambra, 2020). Since COVID-19 outbreak, several studies associated with different pathologies have been performed in orderto find out how damaging this new virus is for the human being. Hereby, in our studies we provided a list of human genes that could be particularly affected as a consequence oftheir molecular similarities with viral genes, not only belonging to SARS-CoV-2 but also to SARS andMERS. The malfunction of these genes has been associated with different human pathologies and is in continuous increase. Patients infected with COVID-19 typically present fever and respiratory symptoms. Nevertheless, it has been reported an increased risk for complications of hypertension, congestive heart failure, and atherosclerosis conducting to anincreased presence of cardiovascular comorbidities (Clerkin et al., 2020; Li et al., 2020; Zheng et al., 2020). Also, some patients have experimented gastrointestinal manifestations (Wong et al., 2020), neurologic complications (Bridwell et al., 2020; Dugue et al., 2020), and complications associated with the endocrine and urogenital systems, among others (Wang et al., 2020; Wu et al., 2020). Diseases and collateral effects caused by COVID-19 infections could be a consequence of the malfunction of the genes listed in our work. Therefore, they should be considered to be incorporated into susceptibility population studies for respiratory viral infections. Hereby, these results lay the groundwork for further research in the field of human genetics associated with the new viral infection, COVID-19, caused by SARS-CoV-2 and for the development of antiviral preventive methods.

## 5. Conclusions

In our study, we described the main factors that shape CUB in SARS-CoV-2, SARS and MERS in comparison with highly expressed genes in human lung tissue and revealed matching features with human genes that could have favoured the virus for an incremented pathogenesis. Furthermore, we provided a list of candidate human genes that could be involved in the viral infection and had not been described yet which could be the key for explaining collateral effects and the human susceptibility to viral infectionsandshould be considered to be incorporated into genetic population studies.

## 6. Declarations

### 6.3. Authors’ contributions

L. K wrote and revised the manuscript L.M. designed the study, performed the bioinformatics analysis, wrote and revised the manuscript.

## Legends to supplementary files

**Supplementary file 2:** Phylogenetic tree using 118 virus genomes including the references SARS-CoV-2, SARS, and MERS and related viruses belonging to non-human host as described in Supplementary file 1. The clusters were virus grouped with SARS-CoV-2, SARS, and MERSare highlighted. The accession id is followed by the host the sample was isolated from or the county in the case of different isolation of SARS-CoV-2.

**Supplementary file 3:** Codon pair bias plots and correlation of viral genes against highly expressed human genes in lungs tissues according to the fold-change between the expression level in lung and the tissue with second-highest expression level according to “Human Protein Atlas” (https://www.proteinatlas.org/humanproteome/tissue/lung). P1: KT444582, P2: KY417146, P3: KY417150, P4: MG772934, P5: MN996532, P6: C869678, P7: MF593268, P8: FJ938064, P9: FJ938066, P10: KX432213, P11: Y572035, P12: AY686864, P13: MK679660, P14: NC_039207, P15: NC_038294, P16: T040336, P17: MT040333, P18: MT126808, P19: NC_045512, P20: MT066156, P21: T263074, P22: NC_004718, P23: MT198652, P24: MT233519, P25: MT233523, P26: T118835, P27: MT233526, P28: MT259235, P29: MT263435.

**Supplementary file 4:** Heatmaps clustering using a hierarchical method of viral genes based on the measures of molecular and codon usage patterns. The viruses whose genes were included here were SARS-Cov-2, SARS, and MERS and the closest viruses to the common nodes in the phylogenetic tree of supplementary file 2. Civets SARS (AY686864), bats MERS (KC869678), bats SARS (KY417150), bats SARS-CoV-2 (MN996532), pangolins SARS-CoV-2 (MT040336), human SARS (NC_004718), human MERS (NC_038294), hedgehogs MERS (NC_039207) and the human SARS-CoV-2 (NC_045512).

**Supplementary file 5:** Heatmap of clusters (1 to 4) using a hierarchical method of viral genes forSARS (NC_004718) of the human host and human genes based on the molecular features. CPB correlation is included in the left for each cluster relating the CPB of human genes (horizontal axis) and CPB of the viral genes (vertical axis).

**Supplementary file 6:** Heatmap of clusters (1 to 6) using a hierarchical method of viral genes forSARS (NC_038294) of the human host and human genes based on the molecular features. CPB correlation is included in the left for each cluster relating the CPB of human genes (horizontal axis) and CPB of the viral genes (vertical axis).

