## Supplementary material for "Molecular features similarities between SARS-CoV-2, SARS, MERS and key human genes could favour the viral infections and trigger collateral effects": Supplementary_file_2.pdf

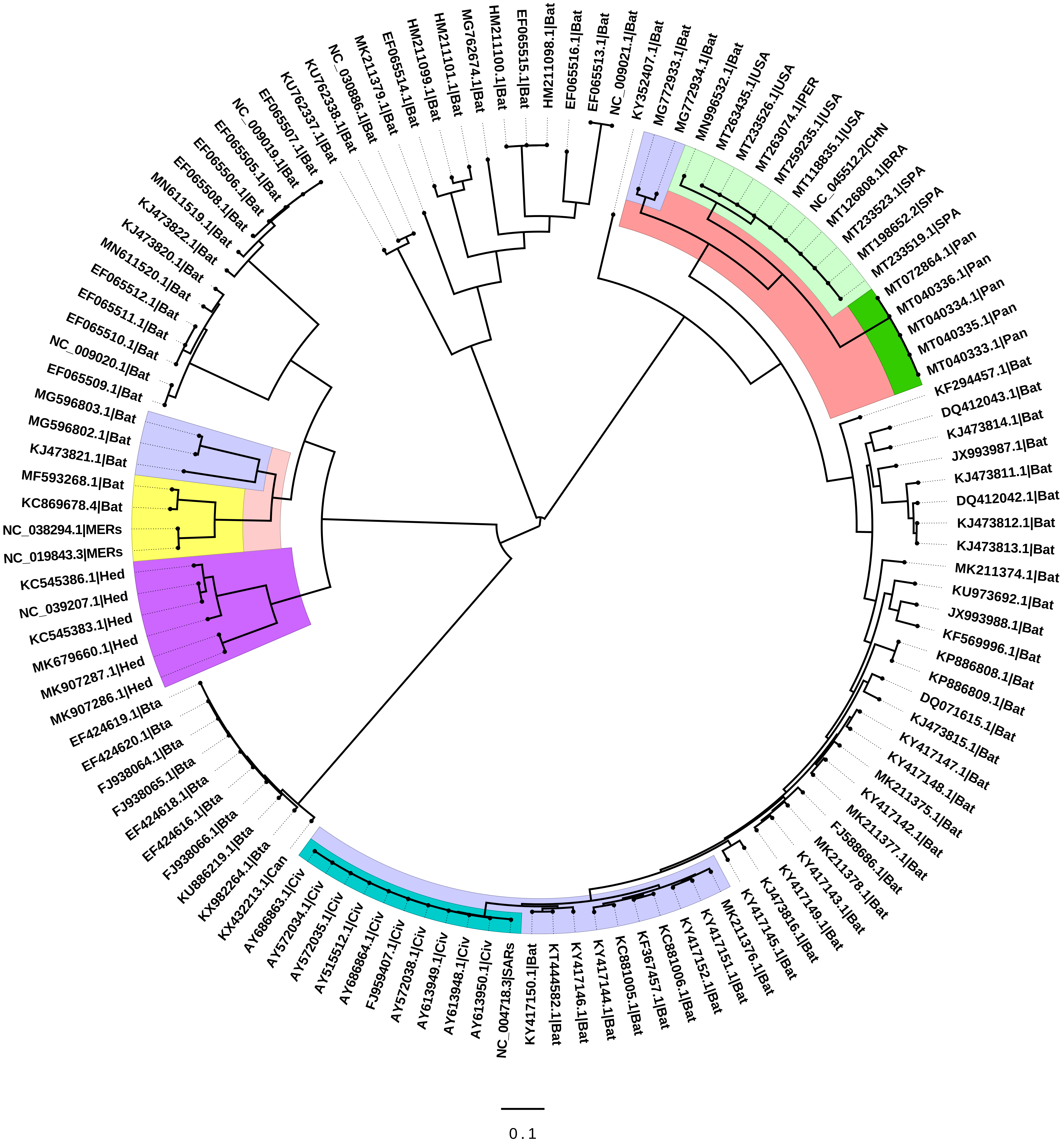


Supplementary file 2: Phylogenetic tree using 118 virus genomes including the references SARS-CoV-2, SARS, and MERS and related viruses belonging to non-human host as described in Supplementary file 1. The clusters were virus grouped with SARS-CoV-2, SARS, and MERS are highlighted. The accession id is followed by the host that the sample was isolated from or the country in the case of different isolations of SARS-CoV-2. Civ: civets, Bta: *Bos taurus*, Hed: hedgehogs, Pan: pangolins.
