## Supplementary material for "Molecular features similarities between SARS-CoV-2, SARS, MERS and key human genes could favour the viral infections and trigger collateral effects": Supplementary_file_3.pdf

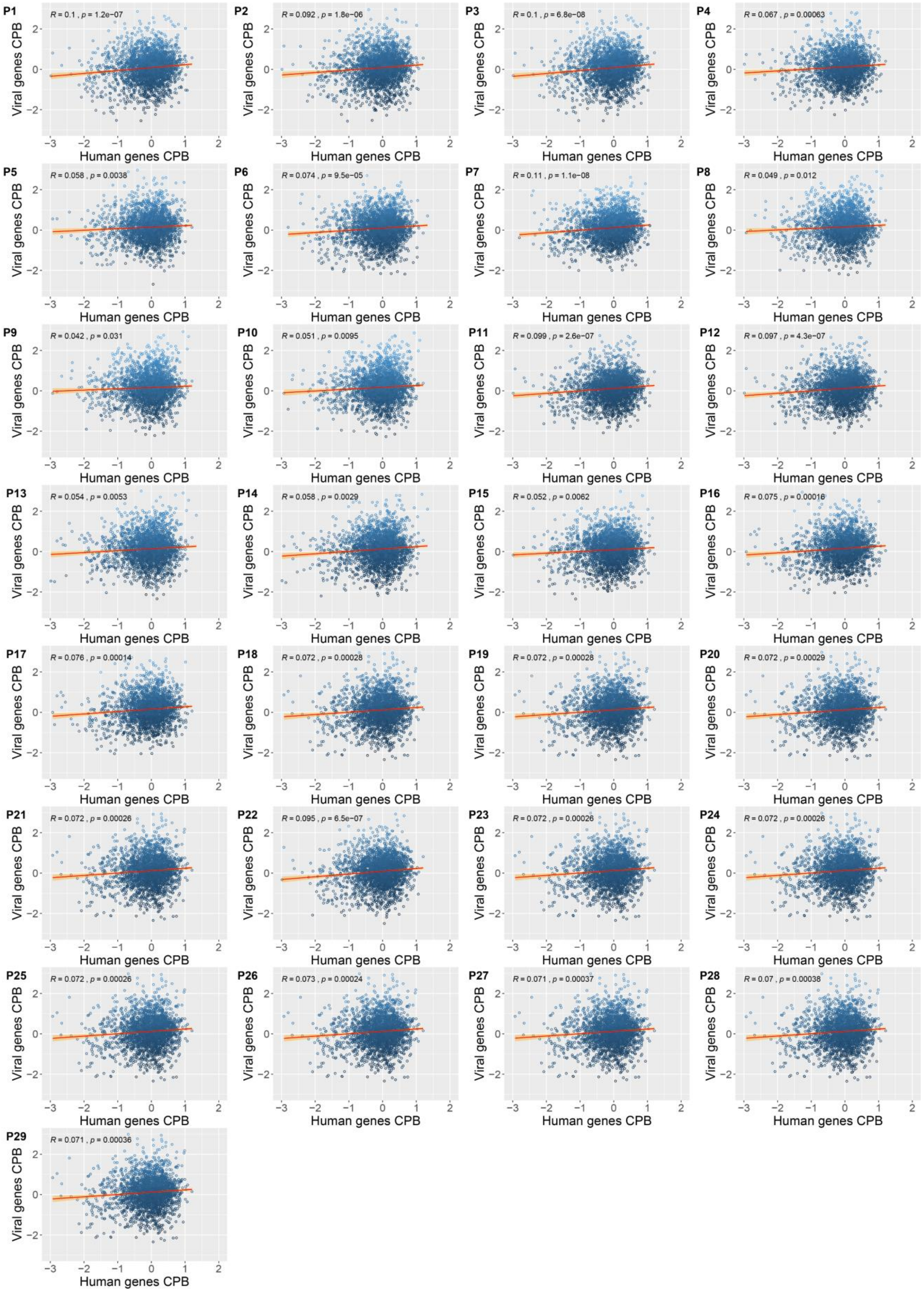


**Supplementary file 3:** Codon pair bias plots and correlation of viral genes against highly expressed human genes in lungs tissues according to the fold-change between the expression level in lung and the tissue with second highest expression level according to "Human Protein Atlas" (https://www.proteinatlas.org/humanproteome/tissue/lung). P1: KT444582, P2: KY417146, P3: KY417150, P4: MG772934, P5: MN996532, P6: C869678, P7: MF593268, P8: FJ938064, P9: FJ938066, P10: KX432213, P11: Y572035, P12: AY686864, P13: MK679660, P14: NC_039207, P15: NC_038294, P16: T040336, P17: MT040333, P18: MT126808, P19: NC_045512, P20: MT066156, P21: T263074, P22: NC_004718, P23: MT198652, P24: MT233519, P25: MT233523, P26: T118835, P27: MT233526, P28: MT259235, P29: MT263435.
