## Supplementary material for "Molecular features similarities between SARS-CoV-2, SARS, MERS and key human genes could favour the viral infections and trigger collateral effects": Supplementary_file_4.pdf

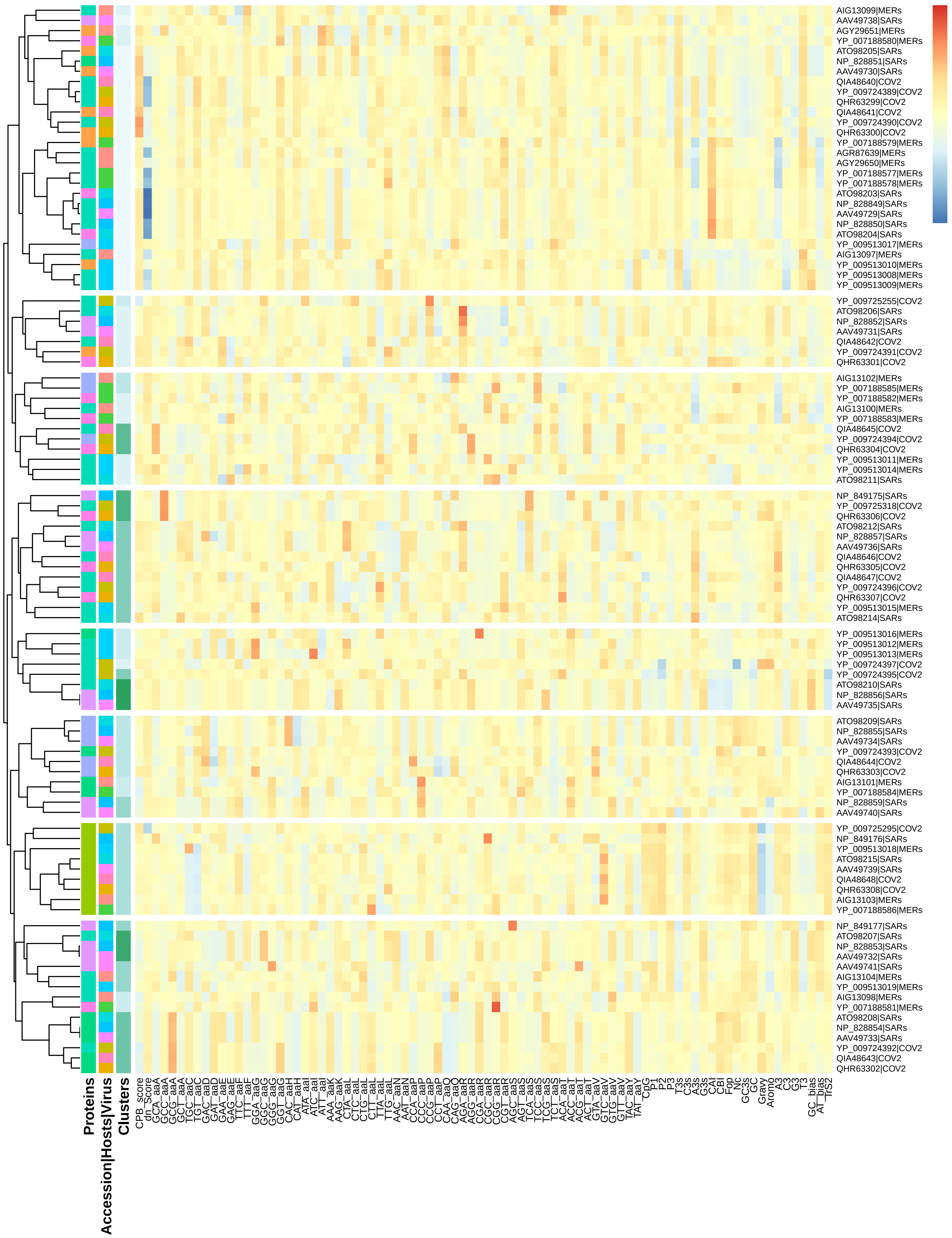


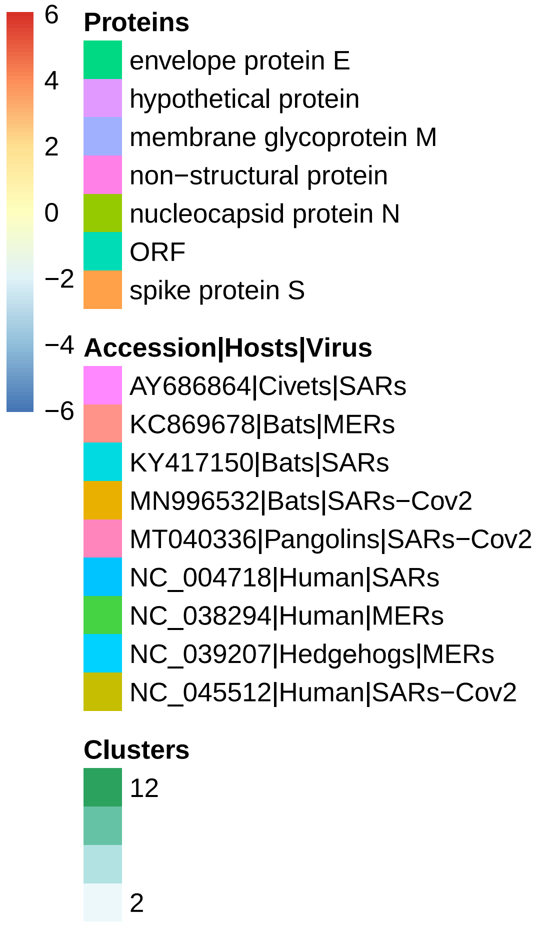


**Supplementary file 4**: Heatmaps clustering using hierarchical method of viral genes based on the measures of molecular and codon usage patterns. The viruses whose genes were included here were SARS-CoV-2, SARS and MERS and the closest viruses to the common nodes in the phylogenetic tree of supplementary file 2. Civets SARS (AY686864), bats MERS (KC869678), bats SARS (KY417150), bats SARS-CoV-2 (MN996532), pangolins SARS-CoV-2 (MT040336), human SARS (NC_004718), human MERS (NC_038294),hedgehogs MERS (NC_039207) and the human SARS-CoV-2 (NC_045512).
