## Supplementary material for "Molecular features similarities between SARS-CoV-2, SARS, MERS and key human genes could favour the viral infections and trigger collateral effects": Supplementary_file_6.pdf

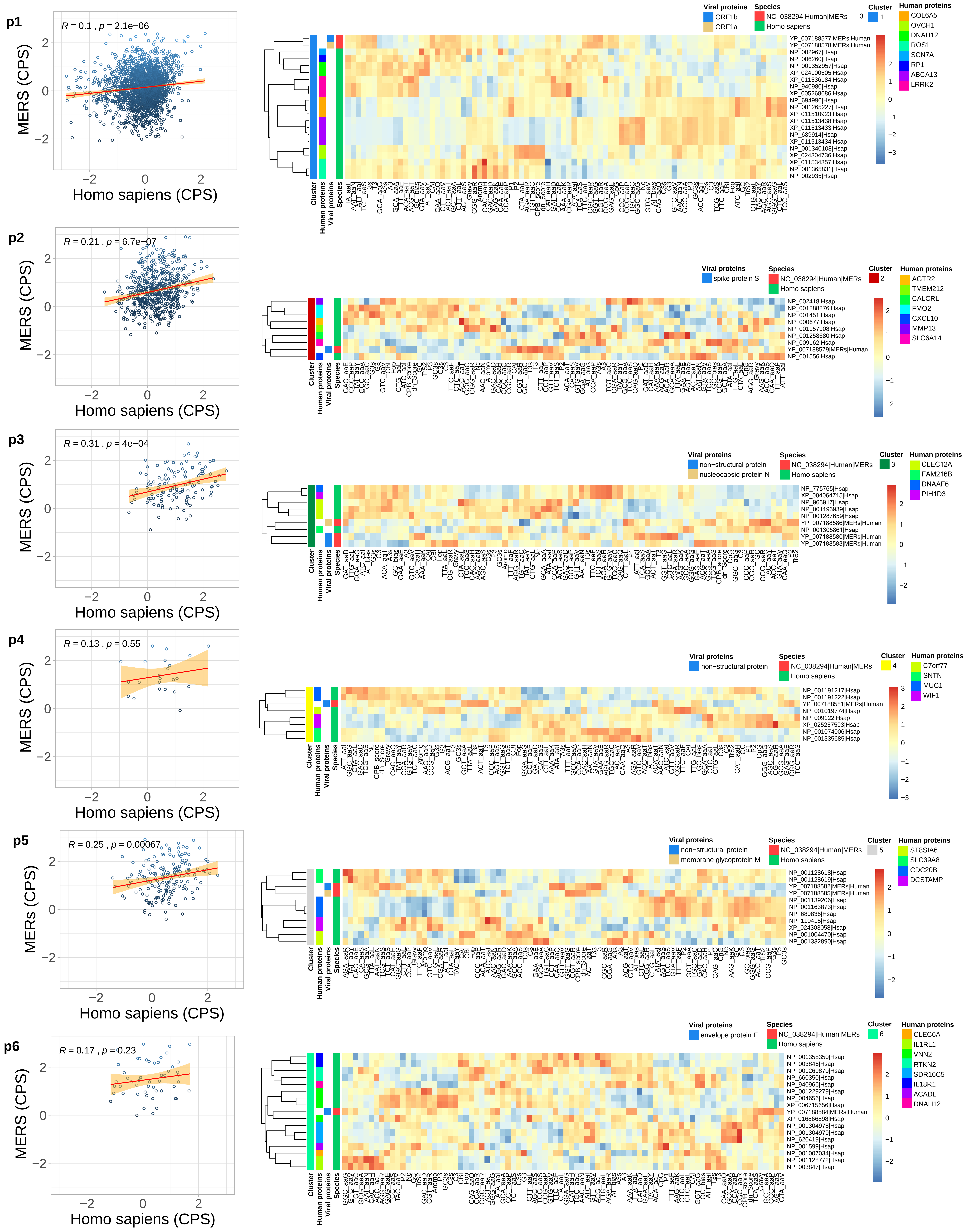


**Supplementary file 6:** Heatmap of clusters (1 to 6) using hierarchical method of viral genes for SARS (NC_038294) of human host and human genes based on the molecular features. CPB correlation is included in the left for each cluster relating the CPB of human genes (horizontal axis) and CPB of the viral genes (vertical axis).
